# Single-cell RNA sequencing identifies KSHV-infected Mesenchymal stem cell subpopulations precursors of Kaposi’s Sarcoma

**DOI:** 10.1101/2023.02.28.530387

**Authors:** Ezequiel Lacunza, Anuj Ahuja, Omar A. Coso, Martin Abba, Enrique Mesri, Julian Naipauer

**Author notes:** Contributed equally (Ezequiel Lacunza, Anuj Ahuja). Deceased (Enrique Mesri).

## Abstract

Kaposi’s sarcoma (KS) may derive from Kaposi’s Sarcoma Herpesvirus (KSHV)-infected human Mesenchymal Stem Cells (hMSCs) migrating into an inflammatory and angiogenic site, enhancing KS initiation. KSHV infection of primary hMSCs uncovers specific cell subpopulations, mechanisms, and conditions involved in the first steps of the KSHV-induced transformation and reprogramming process toward KS progenitor cells. The pro-angiogenic environmental conditions allowed KSHV to reprogram hMSCs closer to KS gene expression profiles and points to KSHV infection as an important factor inducing the Mesenchymal-to-Endothelial (MEndT) transition of hMSC. Single-cell RNA-sequencing analysis revealed a subpopulation of infected cells growing in a pro-angiogenic environment with both viral and host oncogenes upregulation. This highlights the importance of this condition in boosting the KSHV-induced transformation and reprogramming of hMSC towards MEndT and closer to KS gene expression profiles, reinforcing the notion of these cell subpopulations as KS precursors.

## INTRODUCTION

Kaposi’s Sarcoma (KS) is an angioproliferative disease caused by the Human Herpesvirus 8 (HHV-8) or Kaposi’s sarcoma Herpesvirus (KSHV)[1]. Cytokine production and angiogenesis have been shown to contribute to KS pathogenesis [2]; high levels of IL-6, TNF-α, and IL-10 were identified in sera of KS patients[3], and KS tumors have shown elevated levels of IL-6, IL-10, and IFN-γ[4]. The precursor cell for this sarcoma is still under debate because spindle cells in KS lesions express different markers, including endothelial, mesenchymal, and macrophage markers[5]. In this sense, the importance of Mesenchymal Stem Cells (MSC) in KSHV-induced tumorigenesis and transformation, including the increased expression of a large spectrum of chemokines, cytokines, and their receptors after KSHV infection of human MSC (hMSC), has been shown[6–9]. Moreover, KSHV infection of oral mesenchymal stem cells induces mesenchymal-to-endothelial transition (MEndT) and enhances angiogenic factors expression[7]. A study of clinical Kaposi sarcoma specimens has provided evidence for MEndT in Kaposi sarcoma, showing that KS samples are positive for CD105 (Endoglin), CD34, COX-2, VEGF, a-SMA, and c-Kit[10]. Wang et al. recently showed that KSHV infection of periodontal ligament stem cells (PDLSCs) significantly upregulated the expression of several chemokine receptors and promoted MSC migration and settlement in the wound sites[11]. Chen et al. showed that KSHV infection of MSCs initiates an incomplete MEndT process and generates hybrid mesenchymal/endothelial (M/E) state cells and that KS lesions contained a large number of tumor cells with a M/E state[12]. We have previously shown that MSC culture conditions favor viral production with decreased proliferation of infected cells. In contrast, in pro-angiogenic culture, conditions are more permissive for enabling the proliferation of productively infected hMSC cultures. Only KS-like pro-angiogenic conditions induced tumorigenesis in mouse bone marrow-derived MSC infected with KSHV[13].

These data together reinforced the hypothesis that KS may derive from KSHV-infected MSC migrating into an inflammatory and angiogenic site enhancing the transformation and reprogramming capacity of KSHV. However, the cell progenitor subpopulation, defining markers, mechanism, and conditions for KSHV-induced transformation and reprogramming of hMSC toward KS-precursors are not fully understood [14]. Here we carry out KSHV infection of primary bone marrow-derived hMSCs under different environmental conditions. After infection, whole and singlecell RNA-sequencing and pathway analysis of the Differentially Expressed Genes (DEGs) were performed. Differentially regulated pathways related to Extracellular Matrix Organization, Cytokine Activity, Cell Proliferation, MAPK cascade, PI3K-Akt, and Angiogenesis were significantly overrepresented in hMSCs infected in KS-like environments indicating the importance of this condition in KSHV-induced reprogramming of hMSC. Cytokine and Angiogenesis arrays in protein extracts showed upregulation of cytokine and angiogenesis-related proteins in KS-like conditions supporting a boosting effect of KS-like conditions in reprogramming hMSCs towards MEndT by KSHV infection. In addition, we performed unsupervised clustering analysis using a KS gene signature from Tso et al.[15] composed of 3589 significant DEGs between KS lesions and control samples and showed that KSHV-infected hMSC grown in KS-like conditions clustered next to KS lesions and away from the normal skin. Moreover, analysis of 340 genes directly dependent on KSHV infection in KS-like pro-angiogenic conditions showed Endothelial Differentiation as the most significant upregulated pathway, pointing again to KSHV infection as an important factor inducing MEndT in hMSC. Finally, single-cell RNA-sequencing analysis of KSV-infected hMSC showed a subpopulation of infected cells growing in a KS-like environment with upregulation of oncogenic viral together with cytokine and angiogenic-related genes. These results show the importance of the pro-angiogenic growth environment in KSHV reprogramming towards endothelial differentiation and transformation of hMSCs and reinforce the notion that this subpopulation of infected cells would be possible KS precursors.

## RESULTS

### RNA-sequencing analysis of de novo infected bone marrow-derived hMSC in different cell culture conditions

To evaluate the impact of different growth conditions in KSHV infection of bone marrow-derived human Mesenchymal Stem Cells (hMSC), we infected these cells with KSHV in different environments. We performed KSHV (KSHVr.219)[16] infection of hMSC in MEM alpha medium, basal MSC growth conditions (MSC-KSHV^MEM^), or KS-like pro-angiogenic conditions (MSC-KSHV^KS^). The cellular mechanisms promoting inflammation, wound repair, and angiogenesis, may promote the development of KS tumors in KSHV-infected individuals (KS-like media). After 72 hours of infection, we extracted RNA to perform a whole RNA-sequencing analysis. Mock-infected cells growing in basal MSC growth conditions were used as uninfected controls (MSC^MEM^). Mock-infected hMSCs growing in KS-like pro-angiogenic conditions (MSC^KS^) were included as an additional control to uncover the effects of the pro-angiogenic environment (Figure 1A). Principal component (PCA) analysis showed that KSHV infection and KS-like growth conditions induced profound changes in the hMSC transcriptome (Figure 1B). We performed different comparisons to analyze the differentially expressed genes (DEGs) across the samples and better understand the gene expression reprogramming effect of KSHV infection (figure 1C). KSHV infection in MEM medium (MSC^MEM^ versus MSC-KSHV^MEM^) results in 441 DEGs, and infection in KS-like pro-angiogenic conditions (MSC^MEM^ versus MSC-KSHV^KS^) results in 1521 DEGs (Figure 1C, comparisons one and two, Table 1) showing the strong effect of the different environments on host gene expression after KSHV infection. We further performed functional enrichment analysis on DEG from the comparisons of MSC^MEM^ versus MSC-KSHV^MEM^ and MSC^MEM^ versus MSC-KSHV^KS^ (Figure 1D). Processes such as extracellular matrix organization, angiogenesis, cell differentiation, cytokine activity, cell proliferation, MAPK, and PI3K signaling were significantly more overrepresented in MSC-KSHV^KS^ (Figure 1D). This further indicates the importance of this pro-angiogenic growth environment in KSHV reprogramming of hMSC towards an increase in cytokine expression regulation and endothelial lineage differentiation.

**Figure 1.**
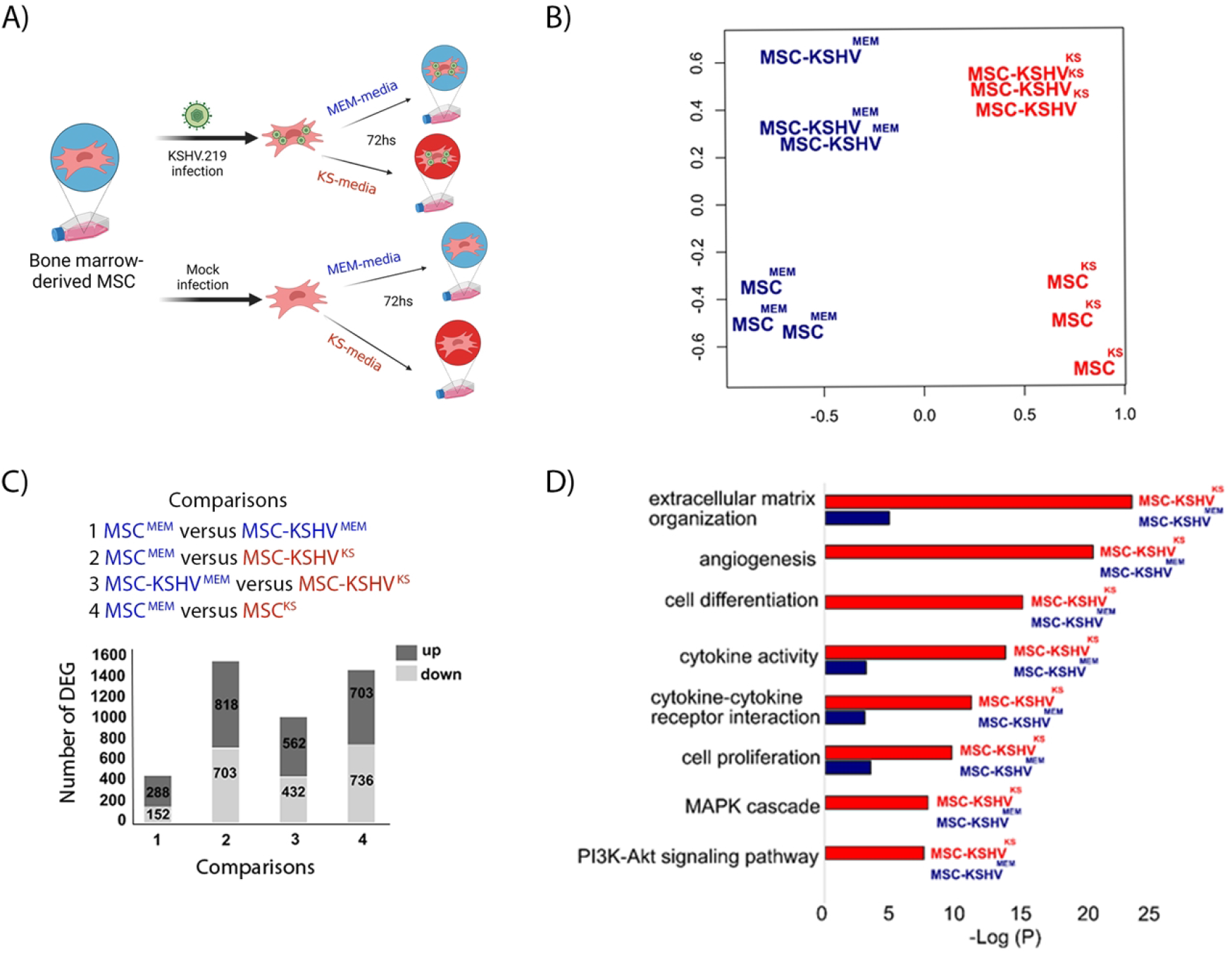
RNA-sequencing analysis of de novo infected bone marrow-derived hMSC in different environments. A) Schematic representation of the sequenced samples. B) Multidimensional scaling plot of the host RNA showing the distance of each sample from each other determined by their leading log Fold Change. C) Key biological comparisons that were detected by RNA-sequencing analysis in triplicates for each sample, and Deferential Expressed Genes (DEGs) in each comparison. D) Functional enrichment analysis based on the DEGs among the two comparisons: MSC^MEM^ versus MSC-KSHV^MEM^, and MSC^MEM^ versus MSC-KSHV^KS^.

### KSHV infection in KS pro-angiogenic environmental conditions reprograms human MSCs to increase Cytokine, Cell proliferation, and Angiogenic markers and leads cells closer to a KS-like cell expression profile

To compare hMSC infected with KSHV in basal MSC growth conditions or KS-like pro-angiogenic conditions, we analyzed the DEGs between the two infected cells MSC-KSHV^MEM^ versus MSC-KSHV^KS^ (comparison three from Figure 1C) and performed pathways analysis of these DEGs (562 up-regulated genes and 432 down-regulated genes, Table 1). We found again Angiogenesis, Extracellular Matrix Organization, MAPK cascade, Cytokine-Cytokine receptor interaction, Inflammatory Response, and Cell Cycle as some of the most differentially regulated pathways (Figure 2A). The heatmap in Figure 2B shows the 140 DEGs from three differentially regulated pathways implicated in KS tumorigenesis (Cytokine Receptor Interaction (yellow), Angiogenesis (red), and Cell Cycle (purple)). Cytokine Receptor Interaction pathway showed CXCL1 (Gro-alpha), CXCL3 (Gro-gama), CXCL6, CXCL8 (IL8), CCL2, CCL7, CCL26, IL1B, and IL33 as the most upregulated genes in MSC-KSHV^KS^ cells. Related to Angiogenesis, we found PECAM1 (CD31), FLT1 (VEGFR1), ROBO4, XDH, NOX5, ESM1, and HGF as upregulated genes. For Cell Proliferation/Cell Cycle we found E2F1, E2F2, E2F8, CDC6, CDC20, CDC45, CDC25A, CCNB2 (Cyclin B2), CDK1, PLK1 and CDKN2D (p19) as upregulated in MSC-KSHV^KS^ cells. Supplementary figure 1 shows the expression of these genes across all the samples (Supplementary Figure 1).

**Figure 2.**
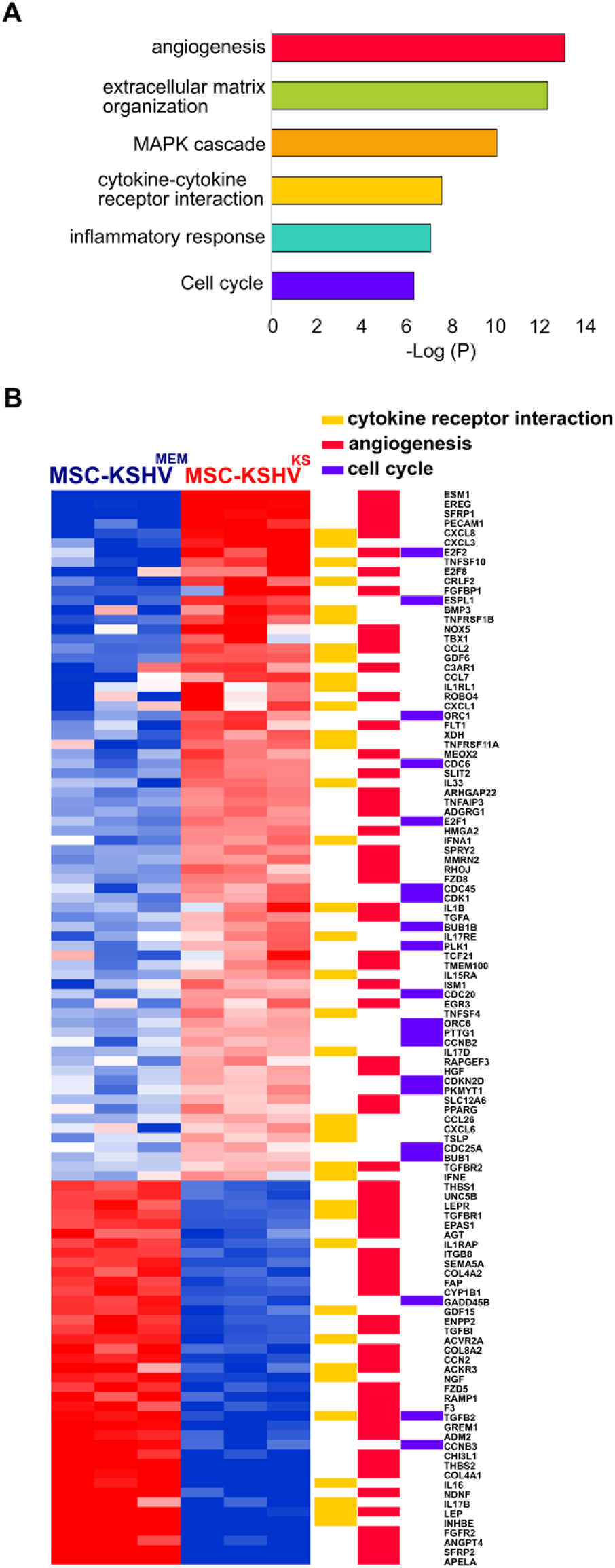
KSHV infection of hMSC in KS environmental conditions showed augment of angiogenesis, cytokine activity and cell proliferation pathways. A) Functional enrichment analysis based on the DEGs among the comparison MSC-KSHV^MEM^ versus MSC-KSHV^KS^. C) Heat map for fold change expression of host RNAs based on analysis of 140 DEGs of three differentially regulated pathways implicated in KS tumorigenesis (Cytokine Receptor Interaction, Angiogenesis and Cell Cycle), upregulated (red) and downregulated (blue) DE genes are shown.

To validate the RNA-sequencing findings, we performed Cytokine and Angiogenesis arrays in protein extracts from MSC-KSHV^MEM^ and MSC-KSHV^KS^ cells, analyzing more than one hundred different cytokines and fifty angiogenesis-related proteins (Figure 3). Interestingly, in the top 20 differentially regulated cytokines, we found upregulation of Endoglin (CD105), basic FGF, CXCL8, iL17, CCL2, Osteopontin, Pentraxin 3, Serpin E1/PAI-1, and Tfr by human MSCs infected in KS-like conditions (Figure 3A). Moreover, the biggest changes in the angiogenesis array showed upregulation of Plasminogen Activator Urokinase (PLAU), or Urokinase-Type Plasminogen Activator (uPA), Pentraxin 3, and Serpin Family E Member 1 (Serpin E1 or PAI1) (Figure 3B). These results support a boosting effect of KS-like conditions in the reprogramming of hMSCs by KSHV infection towards Angiogenesis and Cytokine pathways.

**Figure 3.**
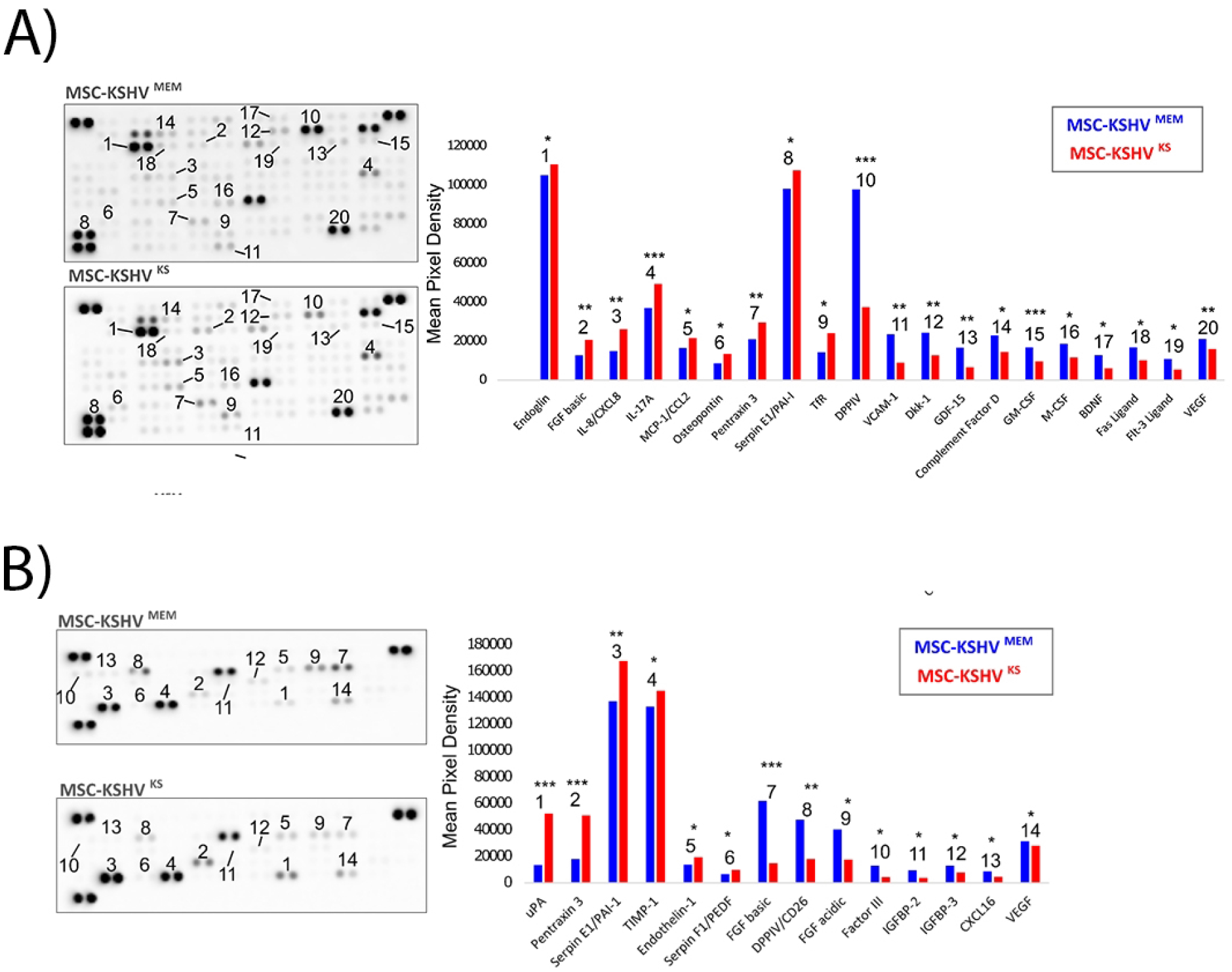
A) Cytokine Array Kit was used to quantify levels of 105 cytokines in MSC-KSHV^MEM^ and MSC-KSHV^KS^ cells pointing to major activation spots corresponding to the top differentially regulated cytokines. B) Angiogenesis Array Kit was used to quantify levels of 55 angiogenesis-related proteins in MSC-KSHV^MEM^ and MSC-KSHV^KS^ cells pointing to major activation spots corresponding to the top differentially regulated proteins.

The results shown in this work indicate that KSHV infection of hMSC in KS-like pro-angiogenic conditions boosts the reprogramming of bone marrow-derived human MSCs to increase cytokine, cell proliferation, and angiogenic markers. As all of these markers are considered KS hallmarks, we decided to perform an analysis using KS signatures from the literature to visualize how KSHV-infected hMSC in different environments correlates with the gene expression profiles of KS lesions. We used a KS signature of 3589 significant DEGs between KS lesions and control samples proposed by Tso et al.[15]. The figure 4A shows the unsupervised clustering - resulting from the gene expression profile of the TSO signature on the samples MSC-KSHV^MEM^ and MSC-KSHV^KS^ from our study, together with samples of KS lesions, contralateral/ipsilateral controls, and normal skin derived from the Lidenge study[17]. MSC-KSHV^KS^ clustered together with all KS samples analyzed, while MSC-KSHV^MEM^ clustered with normal skin and control groups. The same kind of analysis but looking at KSHV gene expression profiles in these same samples showed again that MSC-KSHV^KS^ cluster closer to KS samples than MSC-KSHV^MEM^ (Figure 4B). This analysis indicates that the boosting effect of KS-like conditions of hMSCs by KSHV infection towards Angiogenesis and Cytokine pathways reprograms hMSC closer to KS lesions expression profiles, pointing to these cells as possible KS precursors infected in a pro-angiogenic environment.

**Figure 4.**
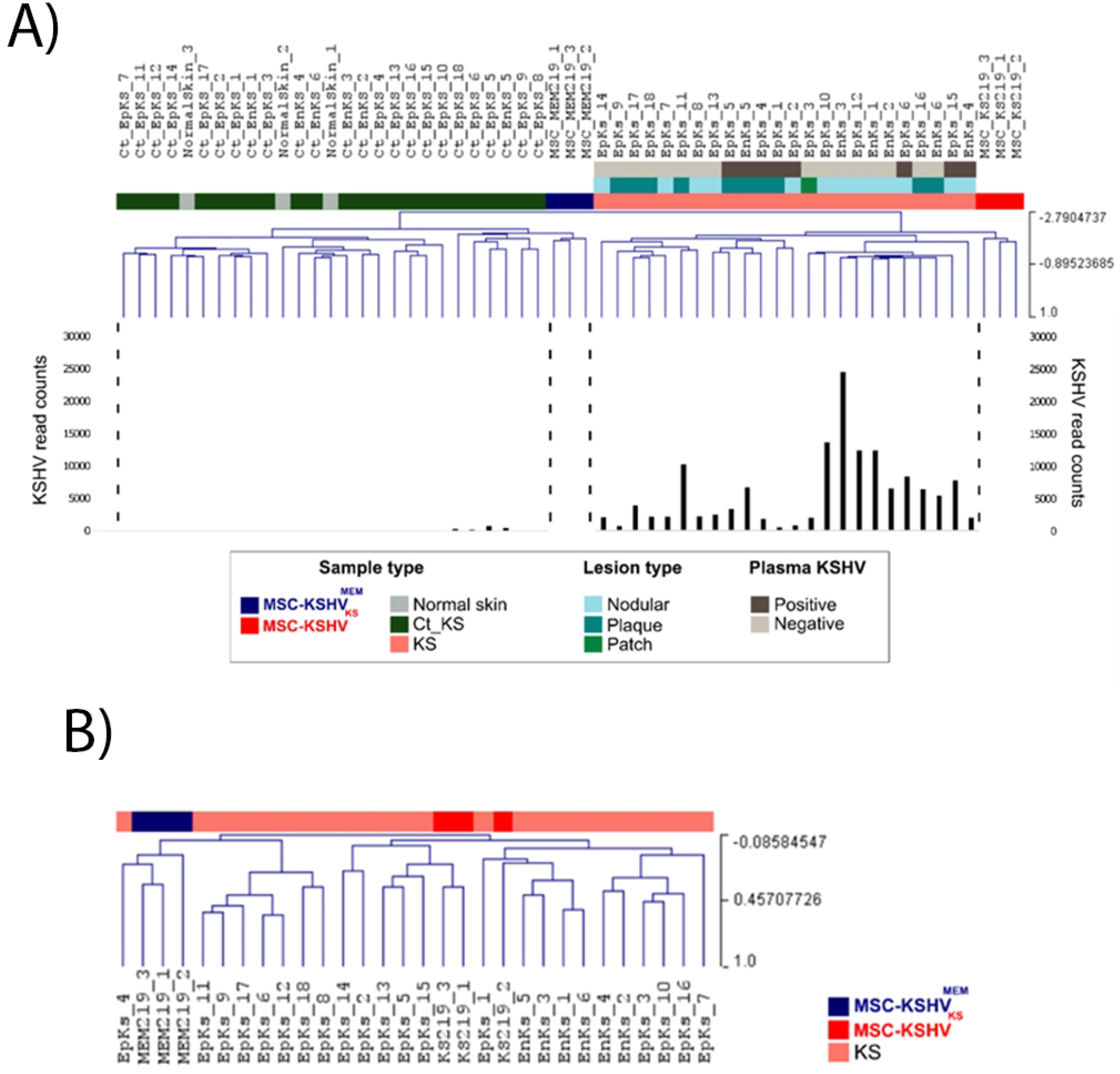
KSHV infection in KS pro-angiogenic environmental conditions reprograms human MSCs in KS-like gene expression profile. A) Unsupervised clustering - resulting from the gene expression profile of a KS signature of 3589 significantly DEGs between KS lesions and control samples-on the samples MSC-KSHV^MEM^ and MSC-KSHV^KS^ from our study, together with 24 samples of KS lesions, 24 contralateral/ipsilateral controls and 3 normal skin derived from the Lidenge study. Lesion type and Plasma KSHV information for each sample were also included. B) Unsupervised clustering - resulting from the KSHV gene expression profile-on the samples MSC-KSHV^MEM^ and MSC-KSHV^KS^ from our study, together with 24 samples of KS lesions from the Lidenge study.

### KSHV infection drives endothelial differentiation of hMSC in the KS-like environment

To understand if the changes in host gene expression towards an increase in Cytokine, Cell Proliferation, and Angiogenic pathways shown in Figures 1, 2, and 3 are induced by KSHV, by the pro-angiogenic environment, or by the combination of both; we analyzed DEGs between hMSC grown in MSC basal or KS-like conditions in the presence or absence of KSHV (Figure 5A). We found 340 genes directly dependent on KSHV infection in KS-like pro-angiogenic conditions; interestingly, pathway analysis of these genes showed Endothelial Differentiation as the most significant upregulated pathway (Figure 5B, Table 1). Among the genes involved in this pathway, ROBO4, IKBKB, XDH, and WNT7B are the most upregulated in MSC-KSHV^KS^ cells, indicating that these genes would be determinants for KSHV-induced reprogramming of hMSC towards Mesenchymal-to-Endothelial transition in the pro-angiogenic environment.

**Figure 5.**
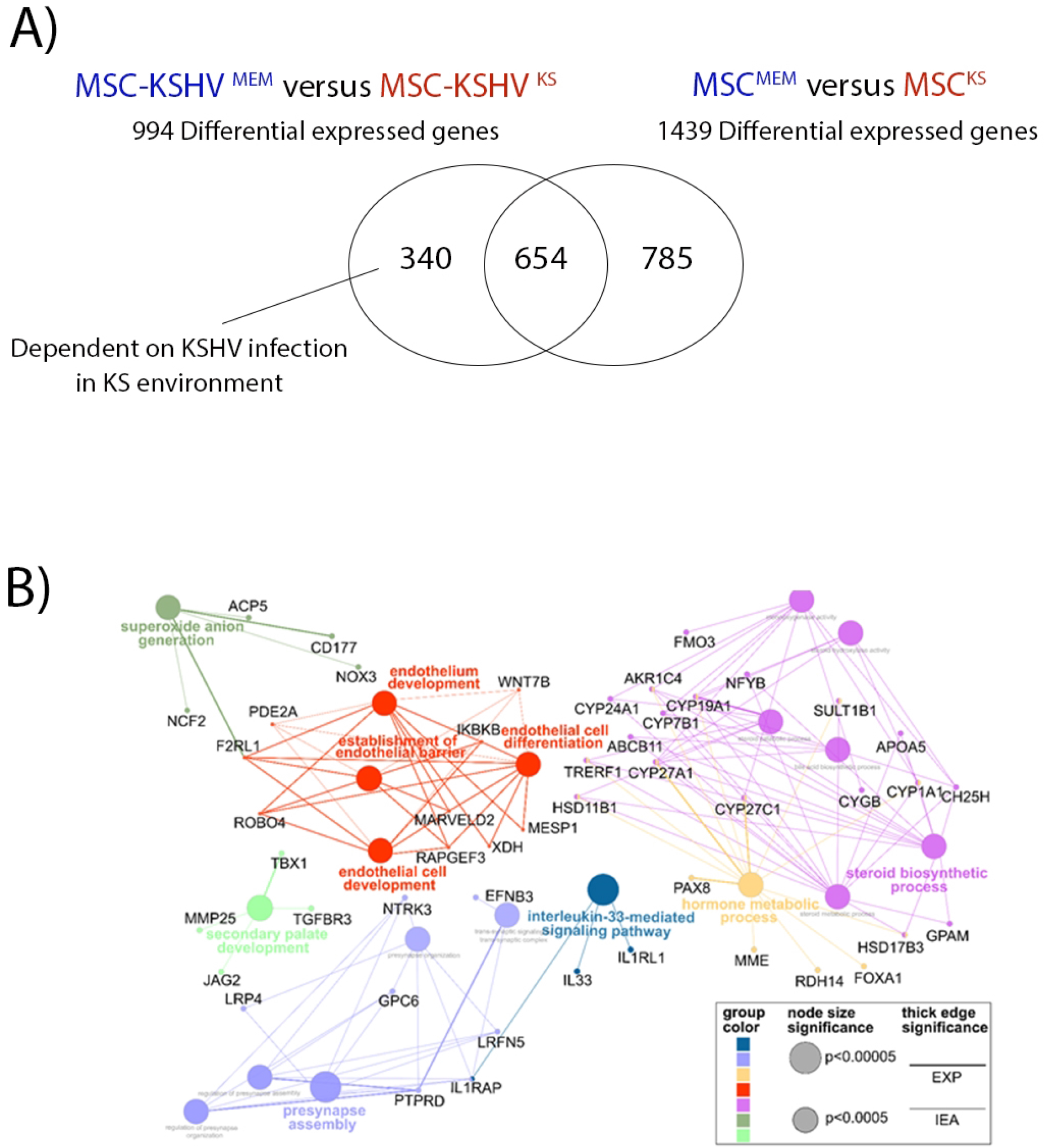
Endothelial differentiation of hMSC in the KS environment is driven by KSHV infection. A) Schematic representation of the comparisons and DEGs found in each one. B) Network analysis of the 340 genes that are directly dependent on KSHV infection in KS-like pro-angiogenic conditions.

### Genome-wide analysis of KSHV transcriptome reveals subtle differences in KSHV gene expression

In the genome-wide analysis of the whole KSHV transcriptome comparing KSHV gene expression between MSC-KSHV^MEM^ and MSC-KSHV^KS^, we found four downregulated (ORF64, K1, K15, and ORF73/LANA) and five upregulated KSHV genes (ORF70, K2/vIL6, ORF57, ORF6, and ORF7) in the KS-like environment (Figure 6A). Supplementary figure 2 shows the expression of most of the latent and lytic KSHV genes in both environments. This paradoxical result goes in the opposite direction from our present and previous results showing that hMSC infection is mostly latent with only 20% of lytic reactivation, pointing to a subpopulation of infected cells as responsible for highly expressing all KSHV genes.

**Figure 6.**
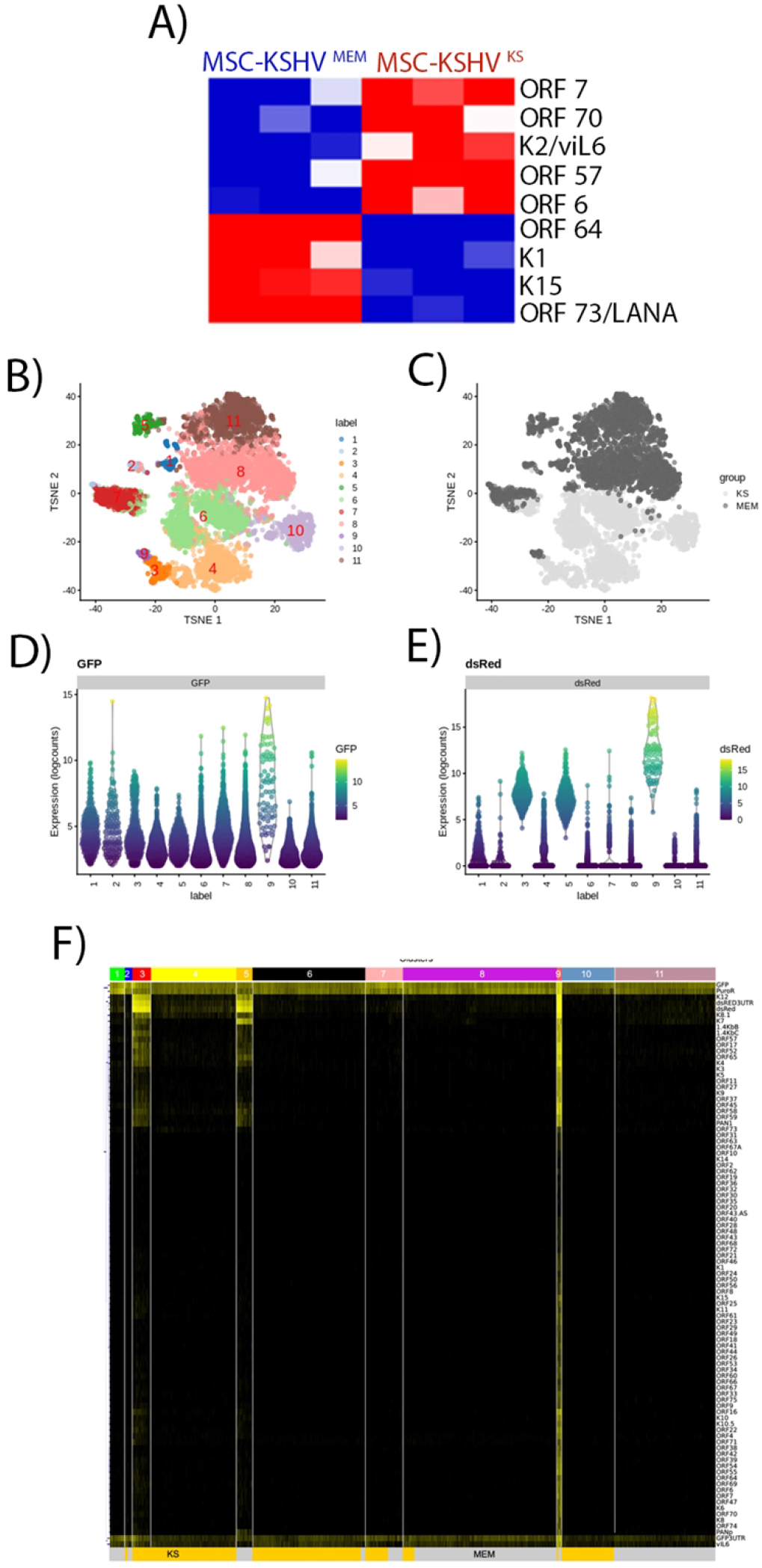
Single-cell RNA-sequencing analysis of human MSC after KSHV infection reveals different subpopulations of infected cells. A) Heat map for fold change expression of KSHV RNAs based on the analysis of DE viral genes between MSC-KSHV^MEM^ and MSC-KSHV^KS^ from the whole RNA-sequencing analysis. B-C) tSNE plot based on viral and host gene expression. Cells are colored according to their cluster identity (B) and the environment in the cells are growing, MEM or KS (C). D) Violin plots showing the expression of GFP (KSHV infection marker) and (E) RFP (KSHV lytic reactivation marker) across the eleven clusters. E) Heat map for fold change expression of KSHV RNAs based on the analysis of the eleven clusters from the single-cell RNA-sequencing analysis.

#### Single-cell RNA-sequencing analysis of human MSC after KSHV infection reveals different populations of infected cells

To reveal whether different infected cell populations contribute to the transcriptomic effects caused by KSHV infection in hMSCs and to uncover the existence of populations of cells highly expressing KSHV genes, we conducted a single-cell RNA-sequencing analysis of bone marrow-derived human MSCs after 72hs of KSHV infection in basal MSC growth or KS-like environment. Over 10,000 high-quality KSHV-infected (GFP-positive) single cells were detected across sequenced samples. High dimensional unsupervised clustering analysis identified eleven cell clusters based on gene expression profiles across the samples in the different environments (Figure 6B, Table 2). Clusters 3, 4, 6, and 10 are enriched in cells (4472 cells) growing in a KS-like environment, while clusters 1, 5, 8, and 11 are enriched in cells (5702 cells) growing in basal MSC media. Clusters 2, 7, and 9 contained a mix of cells growing in both environments (Figure 6C).

We subsequently quantified GFP and RFP expression to uncover latently infected and spontaneous lytic reactivated cells in all the clusters. We found the same average of GFP log counts in all the clusters, pointing to a similar pattern of KSHV latent infection (Figure 6D). Only clusters 3, 5, and 9 showed a high average of RFP log counts; all the cells in these clusters are RFP-positive (Figure 6E), indicating that these cells are undergoing spontaneous lytic reactivation. We found 1549 RFP-positive cells across all the clusters, showing 15% of spontaneous lytic reactivated cells in the total population of infected cells. The heatmap in figure 6F showed KSHV gene expression across the eleven clusters and, as was evidenced by RFP expression, clusters 3, 5, and 9 show the highest levels of expression of KSHV lytic genes (Figure 6F). These results showed that after the novo infection, only a small subpopulation of cells is responsible for the expression of the majority of KSHV genes, with most of the cells only expressing latent and a few lytic genes.

We performed an unbiased analysis of up-modulated host and viral genes in each of the 11 clusters concerning at least 50% of the remaining clusters, with FDR<=0.05 y logFC>=0.58 (Table 2). Interestingly, the most expressed KSHV genes among all the GFP-positive clusters are vIL6, K12, K8.1, and K7. As expected, only the lytic clusters 3, 5, and 9 showed up-modulating KSHV genes compared with the rest of the clusters. Cluster 9, with cells growing in both environments, showed the highest amount of up-modulated viral genes (86 viral genes), indicating that cells in this cluster are undergoing the full lytic program. Importantly, the subpopulation of cells undergoing spontaneous lytic reactivation and growing in the KS-like environment (Cluster 3, 308 cells) showed upregulation of more oncogenic viral genes (including vIRF1, vIRF3, vBcl-2, ORF37/SOX, and vIL6) than the spontaneous lytic reactivated cells growing in basal MSC conditions (Cluster 5, 268 cells) (Figure 7A).

**Figure 7.**
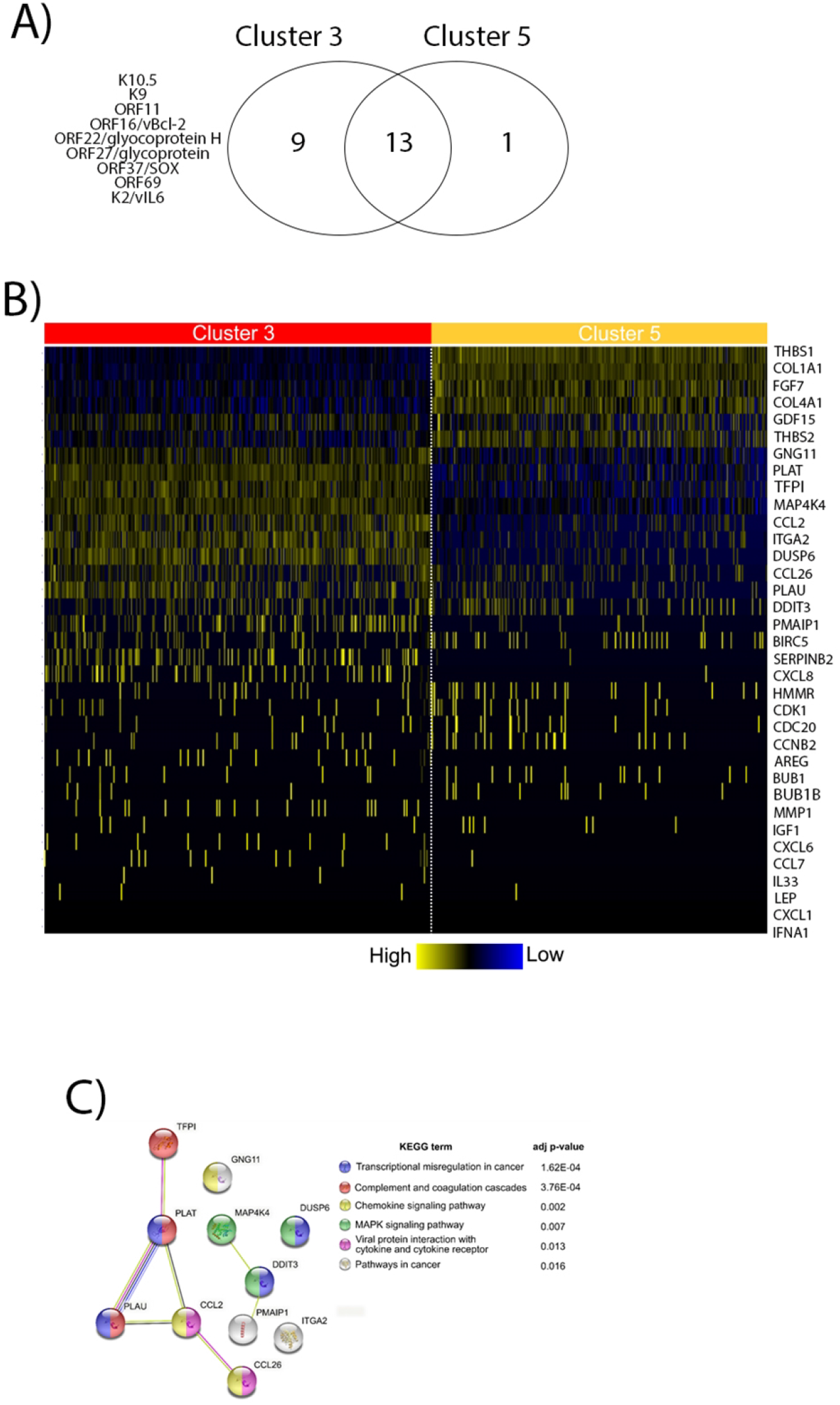
A) Venn diagrams showing upregulated KSHV genes in Cluster 3 and Cluster 5, exclusive KSHV genes for each cluster are highlighted. B) Heat map for fold change expression of 34 KS-related cytokines and angiogenic genes that are differentially regulated between clusters 3 and 5. C) Functional enrichment analysis based on the most upregulated genes in cluster 3.

We analyzed in the single-cell clusters the presence and the expression pattern of the 140 KS-related cytokines and Angiogenic DEGs pathways shown in the heatmap of figure 2B. We found 34 differentially regulated genes between clusters 3 and 5 (Figure 7B). Importantly, the population of cells from cluster 3 showed upregulation of host cytokine-related oncogenes and host endothelial-differentiation markers, indicating that these cells are the only population able to co-express viral oncogenes with host oncogenes; this would allow them to start an oncogenic transformation mechanism in KS-like environment. Pathway analysis of these upregulated genes in cluster 3 showed Transcriptional misregulation in cancer, the Chemokine signaling pathway, and Viral protein interaction with cytokine and cytokine receptors as the most regulated pathways (Figure 7C). One of the up-regulated genes in cluster 3 is PLAU/uPA, the most upregulated angiogenesis protein in the Array shown in Figure 3. These results indicate that after 72 hours of KSHV infection, the different population of infected hMSC expresses different levels of host and viral genes depending on the environmental conditions in which the cells are growing. Pointing to a subpopulation of cells growing in a KS-like environment, with upregulation of oncogenic viral and host genes, as a possible KS progenitor cell subpopulation.

### Single-cell RNA-sequencing analysis of human MSC after KSHV infection and selection

To reveal whether the transcriptomic effects caused by KSHV infection in hMSC at 72hs post-infection are fixed after selection for infected cells and latency establishment, we conducted a single-cell RNA-sequencing analysis of KSHV-infected bone marrow-derived human MSCs after one month of Puromicyn selection in basal MSC or KS-like environment. Over 14,000 high-quality cells were detected and analyzed, importantly all cells analyzed were KSHV-infected (GFP-positive) after selection. High dimensional unsupervised clustering analysis identified nine cell clusters based on gene expression profiles across the samples in the different environments (Figure 8A, Table 3). No clusters with mixed cells growing in the two environments were detected, indicating complete phenotypic differentiation between these infected cells after long-term selection. We detected six clusters of cells growing in the KS environment and only three clusters of cells growing in basal MSC media, indicating more phenotypic cell variability when KSHV infection occurs in a pro-angiogenic KS-like environment (Figure 8B). As expected, all the cells showed GFP expression and the GFP average expression was similar in all the clusters, indicating that all the cells are latently infected with KSHV (Figure 8C). Interestingly, only a few sporadic spontaneous lytic reactivation was seen in the clusters, with clusters 1 and 2 showing the majority of the RFP-positive (lytic reactivated) cells (Figure 8D). The heat map in figure 8E showed all KSHV gene expression over the nine clusters and showed that K12 (Kaposin) and K2 (vIL6) were the most expressed genes all over the clusters. Interestingly, cells in clusters growing in a KS-like environment (clusters 4 and 5) showed upregulation of KSHV genes such as LANA, vFLIP, K8, ORF45, and K1. Again, these results show that after de novo infection and long-term selection, a small subpopulation of cells expressed high levels of KSHV genes and the majority of the cells expressed few viral genes.

**Figure 8.**
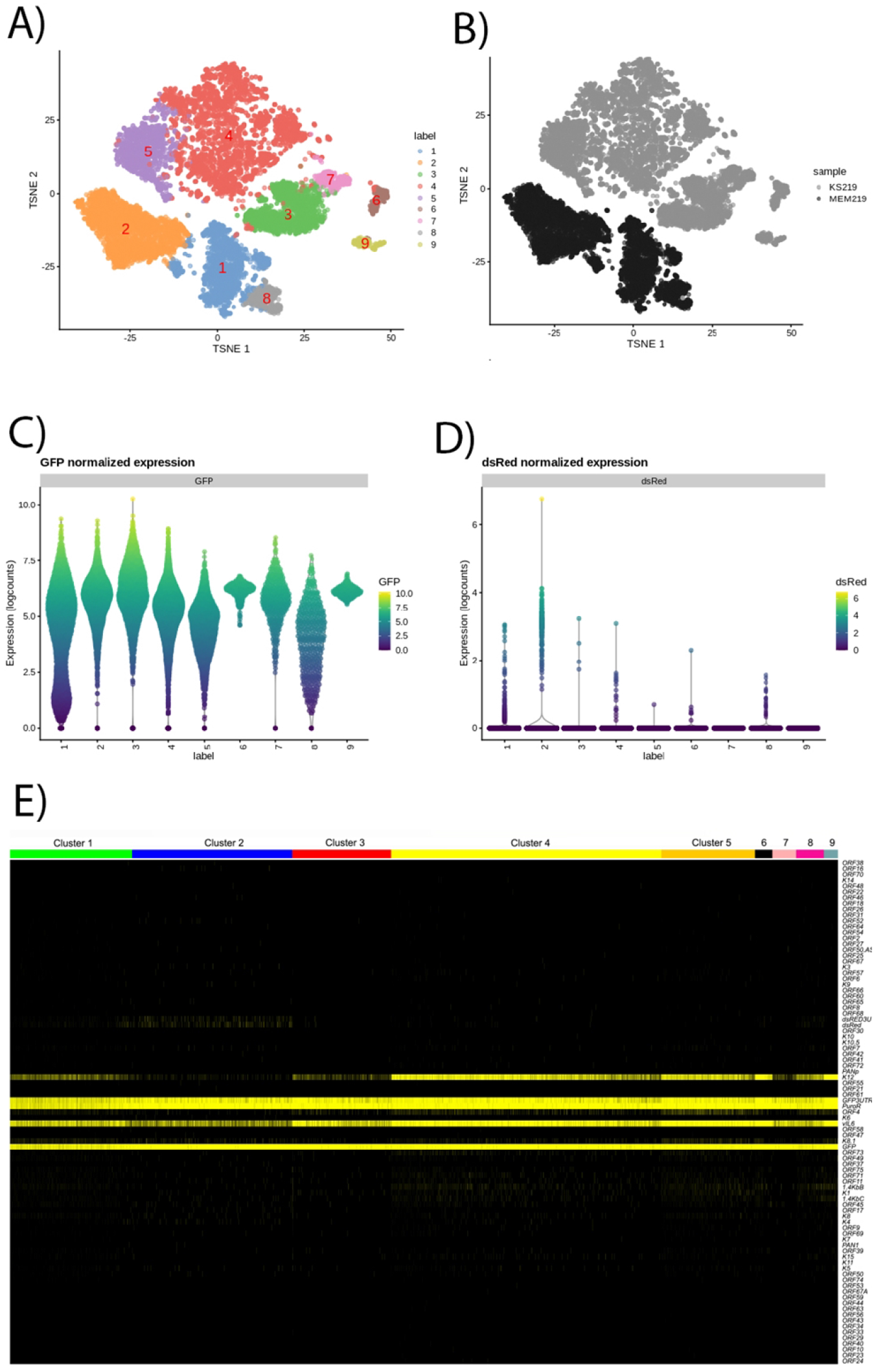
Single-cell RNA-sequencing analysis of human MSC after KSHV infection and selection reveals different subpopulations of infected cells. A-B) tSNE plot based on viral and host gene expression. Cells are colored according to their cluster identity (A) and the environment in the cells are growing, MEM or KS (B). C) Violin plots showing the expression of GFP (KSHV infection marker) and (D) RFP (KSHV lytic reactivation marker) across the eleven clusters. E) Heat map for fold change expression of KSHV RNAs based on the analysis of the eleven clusters from the single-cell RNA-sequencing analysis after long-term selection of infected-cells.

We performed an unbiased analysis of up-modulated host and viral genes in each of the nine clusters concerning at least 50% of the remaining clusters, with FDR<=0.05 y logFC>=0.58 (Table 3). The three clusters detected in basal MSC media shared most of the host genes, with Oxidative Phosphorylation and Translation as the most up-modulated pathways. The six clusters of cells growing in the KS-like environment showed upregulation of angiogenic and cytokine-related genes correlated with upregulation of KSHV gene expression. Among these host genes PLAU, PLAT, HMOX1, ANGPTL4, CXCL12, VEGF, and PTX3 are upregulated and correlated with a subpopulation of cells discovered in the analysis from 72hs post-infection (Figure 7). Our analysis point to these genes as important for the KSHV-induced reprogramming effect of hMSC and as possible markers for a population of cells with host and viral oncogene expression.

## DISCUSSION

Kaposi’s sarcoma remains potentially life-threatening for patients with advanced or ART-resistant disease, where systemic therapy is indicated and three FDA-approved agents, including liposomal anthracyclines, are available[18–20]. Despite the effectiveness of these agents, most patients progress and require additional therapy[21]. Great efforts have been made to identify the KS progenitor primary cell type and the environmental conditions leading to sarcomagenesis. An emerging picture of the origin of the KS progenitor cell point to the Endothelial to Mesenchymal transition axis as important to explain the origin of KS [14]. A better understanding of these processes would allow the development of more accurate and physiologically relevant KS models. Here, we found that processes such as extracellular matrix organization, angiogenesis, cell differentiation, cytokine activity, cell proliferation, MAPK, and PI3K signaling were significantly more overrepresented in Human Mesenchymal Stem Cells infected with KSHV and grown in the KS-like pro-angiogenic conditions together with increase secretion of inflammatory cytokines. The cellular mechanisms promoting inflammation, wound repair, and angiogenesis, may promote the development of KS tumors in KSHV-infected individuals (KS-like media). Using a KS gene signature, we showed that KSHV infection in this environment reprograms human MSCs in a closer KS-like gene expression profile than infection in MSC cell culture conditions. For the first time, Single-cell RNA-sequencing analysis of *de novo* KSHV-infected cells revealed populations of cells expressing different amounts of host and viral genes. One such subpopulation of infected cells in a KS-like environment showed upregulation of oncogenic viral and host genes, further demonstrating the need for pro-angiogenic KS-like culture conditions for oncogenic viral gene expression, virus-induced reprogramming towards Mesenchymal-to-Endothelial transition axis and KS progenitor cell transformation initiation.

The whole RNA-sequencing analysis of hMSC infected in two different environments showed that KSHV infection and pro-angiogenic conditions significantly impact the hMSC transcriptome, directed by KSHV gene expression and the plasticity of hMSC growing in an environment rich in angiogenic factors (Figures 1 and 2). This plasticity allowed KSHV to reprogram hMSC to upregulate the expression and secretion of inflammatory cytokines like Endoglin (CD105), basic FGF, CXCL8, IL17, CCL2, Osteopontin, Pentraxin 3 (PTX3), Serpin E1/PAI-1 and Tfr (Figure 3). Importantly, it was shown that cytokines have an important role in KS pathogenesis[7, 22–25]. After KSHV infection in pro-angiogenic conditions, hMSC expresses high levels of PECAM1 (CD31), FLT1 (VEGFR1), ROBO4, XDH, NOX5, ESM1, and HGF implicated in Angiogenesis and endothelial lineage differentiation. These results support the reprogramming effect of KSHV to induce Mesenchymal-to-Endothelial transition in MSCs and the boosting effect of the KS-like environment.

Using a KS gene signature of 1,482 genes, Li et al. found that KSHV-infected oral MSC cluster closer to KS lesions than KSHV-infected endothelial cells HMVECs, LECs, and BECs[7]. We used a KS signature of 3589 genes[15] and RNA-sequencing of 24 KS samples[17] to show that hMSC infected with KSHV in pro-angiogenic conditions recapitulates better the gene expression profile of KS lesions (Figure 4). This highlight the importance of this condition in KSHV reprogramming of hMSC towards endothelial differentiation and transformation and closer to KS expression profiles, reinforcing the importance of these cells and conditions for KSHV-induced transformation.

We have previously shown that de novo KSHV infection of hMSC leads to a latent and lytic infection and showed that lytic reactivation differs depending on the environmental conditions in which hMSCs are infected. With more virus production and lytic reactivation in hMSCs infected in MEM conditions compared with KS-like conditions. Importantly, we showed that hMSC infection is mostly latent with only 20% of lytic reactivation, and qRT-PCR showed no differences in KSHV LANA, RTA, and K8.1 gene expression[13]. Interestingly, correlated to our previous work, this new genomewide analysis of all the KSHV transcripts showed a slight increase in expression of KSHV genes (ORF70, K2/vIL6, ORF57, ORF6, and ORF7) in a pro-angiogenic environment (Figure 6A). However, KSHV-infected hMSC in both environments showed expression of latent and lytic KSHV genes (Supplementary Figure 2). These slight differences in KSHV gene expression and the fact that we observed latent and lytic KSHV gene expression would be due to subpopulations of infected cells expressing different amounts of viral genes, which would be obscure by the analysis of whole populations of cells made by qRT-PCR or whole RNA-sequencing analysis. In this line, our single-cell RNA-sequencing analysis showed the existence of different subpopulations of infected cells based on the amount of KSHV and host gene expression (Figure 6), including different subpopulations of cells that contain spontaneous lytic reactivated cells with only one small of these subpopulations undergoing the full lytic program (Figure 6 and 7). Importantly the small number of cells undergoing the full lytic program also correlated with our previous work, where we showed little virus production in infected hMSC[13]. This is the first study showing single-cell RNA-sequencing data after de novo KSHV infection of KS progenitor cells, showing that a small number of cells undergo spontaneous lytic reactivation, including subpopulations of cells in different environmental conditions and different degree of KSHV lytic gene expression.

Chen et al. recently found that KSHV can promote an intermediate state of Mesenchymal-to-endothelial differentiation of subpopulations of Periodontal ligament stem cells (PDLSCs), correlated with the existence of KSHV-positive spindle cells in Kaposi sarcoma lesions displaying a mesenchymal/endothelial phenotype (Chen et al., 2021). In this line, our whole and single-cell RNA sequencing transcriptomic analysis showed the existence of subpopulations of KSHV-infected hMSC expressing oncogenic viral genes and oncogenic host genes together with endothelial and mesenchymal differentiation markers, pointing to a role of these cells in KS initiation and progression through the mesenchymal-to-endothelial differentiation axis.

## ACKNOWLEDGMENTS

We want to thank Dr. Marta Maria Gaglia for kindly provide the .gtf/gff file to annotate the GFP and RFP transcripts on the recombinant rKSHV219 virus. The Oncogenomics Core Facility at the Sylvester Comprehensive Cancer Center from the University of Miami for performing high-throughput sequencing.

## IN MEMORIAM

Dr. Enrique A. Mesri, an inspiring scientist, a great mentor, and friend, who sadly passed away days before sending the manuscript. For his dedication and contributions to the KSHV and viral oncology field, we will be always in debt to him.

## MATERIALS AND METHODS

### Cell culture

Human MSCs were isolated as described previously[26]. Briefly, BM aspirates (25–50 mL) were purchased from AllCells (Emeryville, CA) under appropriate informed consent and institutional review board approval. The experiments were performed using MSCs in passages 4–7.

### Virus preparation and infection

iSLK-219 cells harboring recombinant KSHV 219 from Don Ganem and Myoung[27] were used for virus preparation. Briefly, infectious viruses of the 219 strain were induced from the respective iSLK cells by treatment with doxycycline and sodium butyrate for 4 days. The culture supernatants were filtered through a 0.45-μm filter and centrifuged at 25,000 rpm for 2 h. The pellet was re-suspended in phosphate-buffered saline (PBS) and aliquot and stored at −70°C as infectious KSHV preparations. Virus infection was performed according to the method used in a previous study, with minor modifications. Human MSCs were seeded 6 × 104 cells per well in 6-well culture plates. After a day of culture, the culture medium was removed, and cells were washed with PBS. The prepared KSHV inoculum, MOI of 8, and 8 μg/ml of Polybrene were mixed and added to the cultured cells. After centrifugation at 700 × g for 60 minutes, the inoculum was removed after 3 hours, and 2 ml of culture medium was added to each well.

### Whole RNA-Sequencing analysis

60000 hMSCs/well were plated in 6 well plates. 24hs later hMSCs were infected with rKSHV219 [16] and cultured in 2 different environments, MEM or KS-like culture conditions. 72hs after KSHV infection, RNA from hMSC was isolated and purified using the RNeasy mini kit (Qiagen). RNA concentration and integrity were measured on an Agilent 2100 Bioanalyzer (Agilent Technologies). Only RNA samples with RNA integrity values (RIN) over 8.0 were considered for subsequent analysis. We performed paired-end sequencing using a NovaSeq platform. The short-read sequences were mapped to the human reference genome (GRCm38.82) by the splice junction aligner TopHat V2.1.0. We employed several R/Bioconductor packages to accurately calculate the gene expression abundance at the whole-genome level using the aligned records (BAM files) and to identify differentially expressed genes between cell lines and tumors. Briefly, the number of reads mapped to each gene is based on the TxDb. Human gene ensembls were counted, reported, and annotated using the Rsamtools, GenomicFeatures, and GenomicAlignments packages. To identify differentially expressed genes between cells, we utilized the DESeq2 test based on the normalized number of counts mapped to each gene. Data integration and visualization of differentially expressed transcripts were done with R/Bioconductor. KSHV transcriptome was analyzed using previous resources and the KSHV 2.0 reference genome[28], while the edgeR test was employed for differential gene expression analysis of KSHV transcripts.

### Kaposi’s Sarcoma gene expression signature profile

Kaposi’s sarcoma KSHV and host RNA-seq profiles were retrieved from the GEO database GSE147704[17] from a previous study and integrated with hMSC-derived samples for further analysis. For unsupervised clustering analysis we applied Pearson correlation with complete linkage method.

### Single-Cell RNA-sequencing analysis

60000 hMSCs/well were plated in 6 well plates. 24hs later hMSCs were infected with rKSHV219[16] and culture in 2 different environments, MEM or KS-like culture conditions. 72hs post-infection or one month after puromycin selection, the cells were washed with PBS and added Trypsin for 5 minutes. Neutralize the trypsin with media recovered and centrifuge to pellet the cells. Take out media, leaving 10ul in order not to lose pellet cells, and resuspend with 70ul of FACS buffer (PBS+1%BSA+5%FBS, filtered). Four human mesenchymal stem cell single-cell suspensions were submitted for 10X Genomics 3’ v3.1 single-cell RNA-Seq. The samples were QC’ed on a Nexcelom Cellometer K2 with ViaStain AOPI Staining Solution under the assay – hMSC. Fluorescent Assay cell counts are summarized below for each sample run. We performed paired-end sequencing using a NovaSeq platform.

### Human Cytokine and Angiogenic Array

R&D Systems’ Proteome Profiler Human XL Cytokine Array Kit (Catalog # ARY022B) and Proteome Profiler Human Angiogenesis Array Kit (Catalog # ARY007) were used to detecting levels of 105 Cytokines and 55 different angiogenic-related proteins in MSC-KSCMEM and MSC-KSHVKS cells, respectively.

### Statistical analysis

The statistical significance of the data was determined using a two-tailed Student’s t-test and 2-way ANOVA for multiple comparisons. A p-value lower than 0.05 was considered significant. Statistical analysis was performed using GraphPad Prism 7. All values were expressed as means ± standard deviation.

